# Molecular Data From Herbarium Specimens Reveal A Potential Extirpation of *Azolla microphylla* From California

**DOI:** 10.64898/2026.10.16.758102

**Authors:** Michael J. Song, Abbey Schedler, Griffin Wagner, Forrest Freund, Isaac L. Maarck, Carrie M. Tribble, Carl J. Rothfels, Pieter Johnson, Jenna T. B. Ekwealor

## Abstract

Previous studies have shown that there may be more species of *Azolla* in California than currently recognized, including a recent introduction of the non-native *Azolla caroliniana*. To better assess the status and species diversity of *Azolla* in California, we sequenced four historical herbarium specimens, some over 100 years old, and increased our sampling of extant populations across seven counties in California. We found that many specimens from California thought to be *Azolla microphylla* are actually *Azolla filiculoides* or members of one of two non-native “*Azolla caroliniana*” clades. We also found that some specimens collected from California from the 1960s through the 1980s fall within an *Azolla microphylla-mexicana* clade. However, there is no evidence of this species from any recent collections even from counties where it was previously found, which points to a potential extirpation of this species from California.

## Introduction

*Azolla* Lam. is a minute, floating aquatic fern in the order Salviniales. There are two major sections that are currently recognized: *Azolla* sect. *Azolla* and *Azolla* sect. *Rhizosperma* (Mezgar et al., 2007). *Azolla* sect. *Rhizosperma* comprises two unmistakable taxa, *Azolla pinnata* and *Azolla nilotica*, whereas the five species of *Azolla* sect. *Azolla* (*A. filiculoides, A. rubra, A. caroliniana, A. mexicana,* and *A. microphylla*) are all similar to one another and require the examination of microscopic characters for identification (Smith and Murdock, 2012a,b, Lumpkin, 1993). The range of *A. nilotica* is limited to Africa and *A. rubra* to Australia and New Zealand, but all of the other species are now widespread and many are considered noxious weeds (Kay and Hoyle, 2001).

Currently, most taxonomic and floristic treatments recognize two species of *Azolla* as occurring in California: *Azolla filiculoides* Lam. (Smith and Murdock, 2012a) and *Azolla microphylla* Kaulf. (Smith and Murdock, 2012b), the latter of which is considered synonymous with *Azolla mexicana* C. Presl. The morphological distinctions between these two taxa are few and microscopic: *Azolla filiculoides* has leaf lobes with a band of cells more than four cells wide and male spore mass (massula) barbs (glochidia) lacking partitions, whereas *Azolla microphylla* has leaf lobes with a band of cells only 1–2 cells wide and male spore mass barbs with 1–3 partitions (Smith and Murdock, 2012a,b, Lumpkin, 1993). Of these characters, those associated with the reproductive morphology are generally thought to be more reliable for identifying *Azolla,* including glochidia septation and megaspore features like presence of angular bumps, pittedness, and amount of filosum cover (Lumpkin, 1993). However, many *Azolla* individuals are sterile, and Evrard and Van Hove (2004) noted that glochidia septation may not be a reliable character.

Recent studies have found that regional *Azolla* diversity is often underrecognized due to morphological similarity. Two distinct clades of “*Azolla cristata*” were found in Indonesia and corresponded to wild and cultivated habitats, respectively (Najibulloh et al., 2025). Similarly, four different species of *Azolla* section *Azolla* were found in Uganda (Lydia et al., 2023). In California, our previous research suggests that there may be up to five species of *Azolla*: the non-native *A. pinnata* and four taxa in *Azolla* section *Azolla*. These include *A. filiculoides*, *A. microphylla* (reported in earlier literature), and two separate clades currently classified as *A. caroliniana* that may have recently been introduced and displaced the native *A. microphylla* (Song et al., 2023, 2025a,b,c). Given this previously underrecognized diversity, we examined historical specimens from the Jepson Herbarium, including those collected by Willis Linn Jepson, and expanded our sampling of extant California populations to investigate changes in *Azolla* diversity over time.

Our study addresses three questions. First, can vegetative or reproductive morphological traits be used to reliably identify *Azolla*? Second, can herbarium specimens yield phylogenetically informative data for *Azolla*? And third, do historical and contemporary California *Azolla* collections represent the same taxa, or is there evidence of a recent cryptic invasion?

## Methods

### Sampling

We used the Consortium of California Herbaria (CCH; CCH Portal, 2026) to identify historical *Azolla* specimens with fertile material and that had been determined as *A. microphylla* by Alan Smith, one of the authors of the Jepson Manual’s treatments of *Azolla* (Smith and Murdock, 2012a,b). We selected four specimens from the Jepson Herbarium based on them being the three oldest specimens identified by Smith as *A. microphylla* based on reproductive morphology that had sufficient material available for sampling, as well as one specimen from the 1960s, a period during which many other collections of *A. microphylla* were made in California:

1. JEPS35135 *Azolla microphylla* Kaulf., by A. Smith, 2006. Collector: Willis L. Jepson 13325. Aug 18, 1928. Irrigation canal north of Dos Palos, Merced County, CA. Mega and microspores present. Septate glochidia.
2. JEPS59976 *Azolla microphylla* Kaulf., by A. Smith, 2006. Originally *Azolla filiculoides* Lam. Collector: Hilda W. Grinnell. Oct 9, 1933. 1 mile north of Joe Walker Mine (South side Basin Creek); Walker Basin, Kern County, CA. Sporangia present.
3. JEPS27903 *Azolla microphylla* Kaulf., by A. Smith, 2006. Originally *Azolla filiculoides* Lam. Collector: Vernon Huebschwerlen 11. Apr 20, 1962. North bank San Joaquin River at Santa Fe Bridge (ca. 110 yds from bridge), Fresno, CA. Sporangia present.
4. JEPS59977 *Azolla microphylla* Kaulf., by A. Smith, 2006. Originally *Azolla filiculoides* Lam. by Jepson. Collector: Willis L. Jepson 10249. Oct 18, 1923. Lake Merced, San Francisco, CA. Sporangia present.

We also expanded our field sampling of contemporary California *Azolla* populations to determine whether *A. microphylla/mexicana* currently occurs in the state, as it was not detected during our 2021 sampling (Song et al., 2025b,c). We collected *Azolla* from 21 additional sites across seven counties (Table 1): Alameda (Pleasanton Ridge Regional Park, Vallejo Mills, Augustin Bernal Park, Hayward), Mono (Mammoth Lakes), San Joaquin (Mokelumne River Day Use Area), Contra Costa (Briones Regional Park, San Pablo Reservoir, East Bay Municipal Utility District Hampton Staging Area), Glenn (Sacramento National Wildlife Refuge Admin Building and Visitor Contact Station), Mendocino (Hopland), and Shasta (Anderson). All collections were made with required permits or permissions. Specimens were deposited in the Harry D. Thiers Herbarium (SFSU) at San Francisco State University, and sequences were submitted to NCBI GenBank to be made publicly available.

**TABLE 1.**
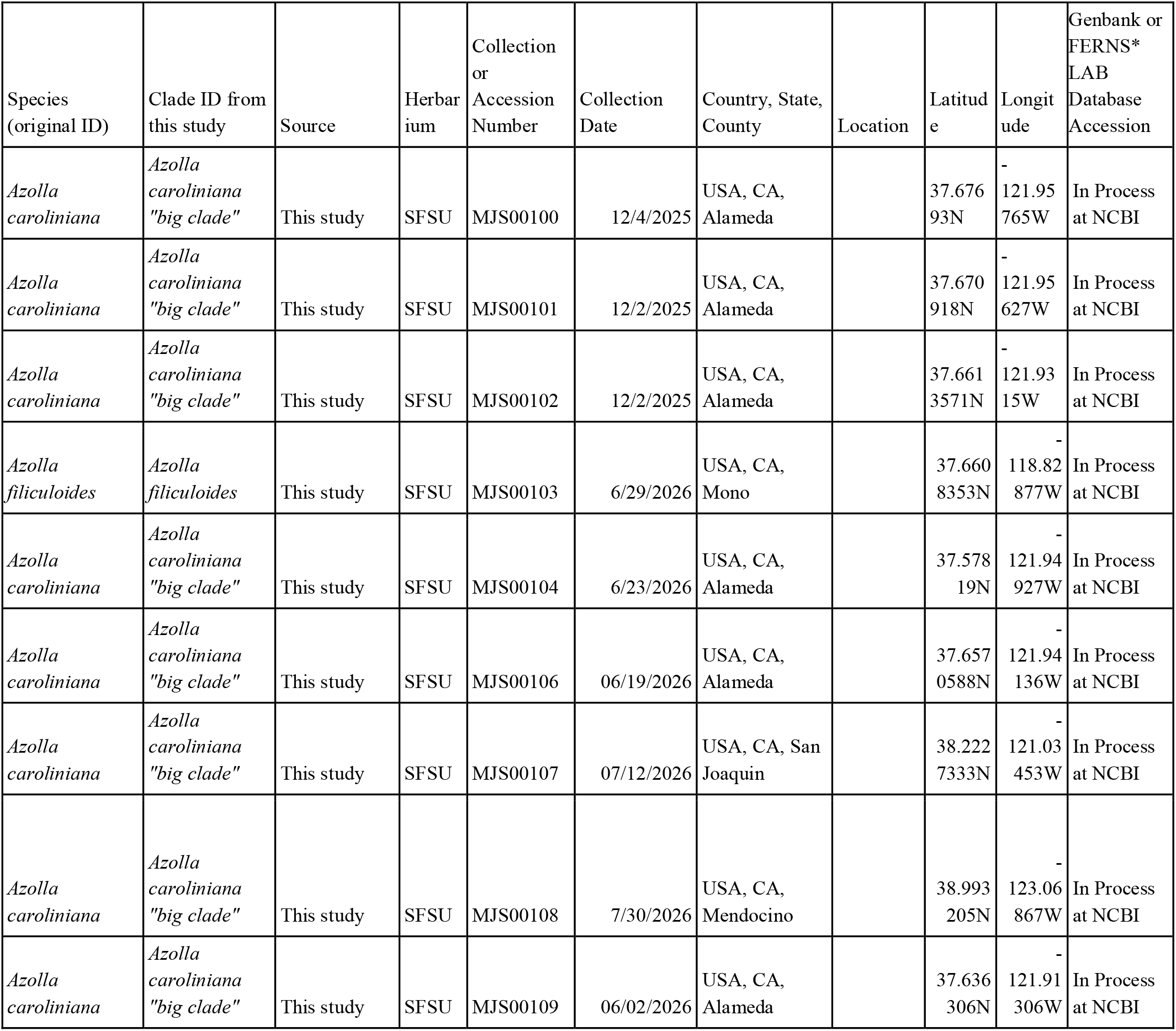
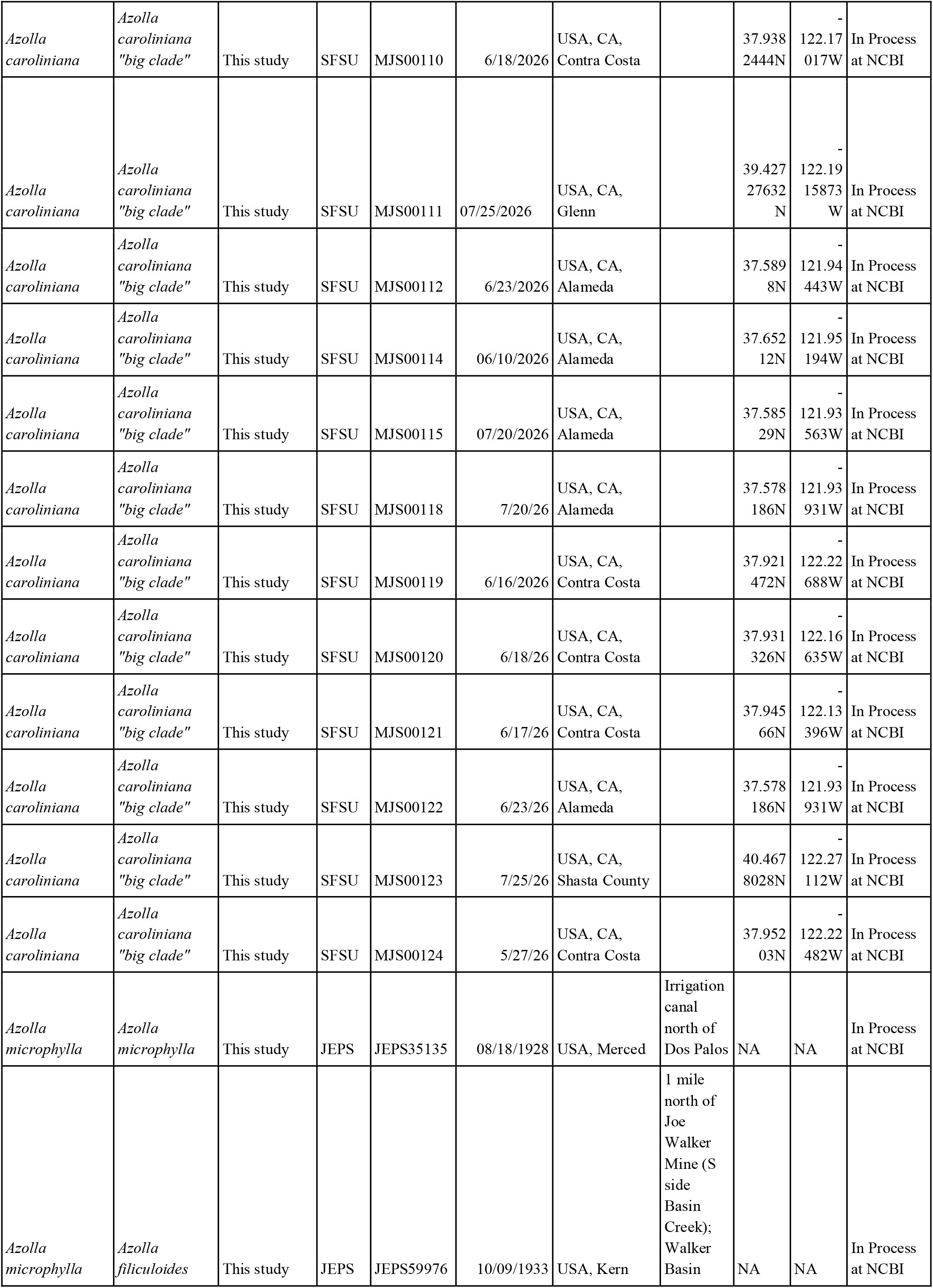
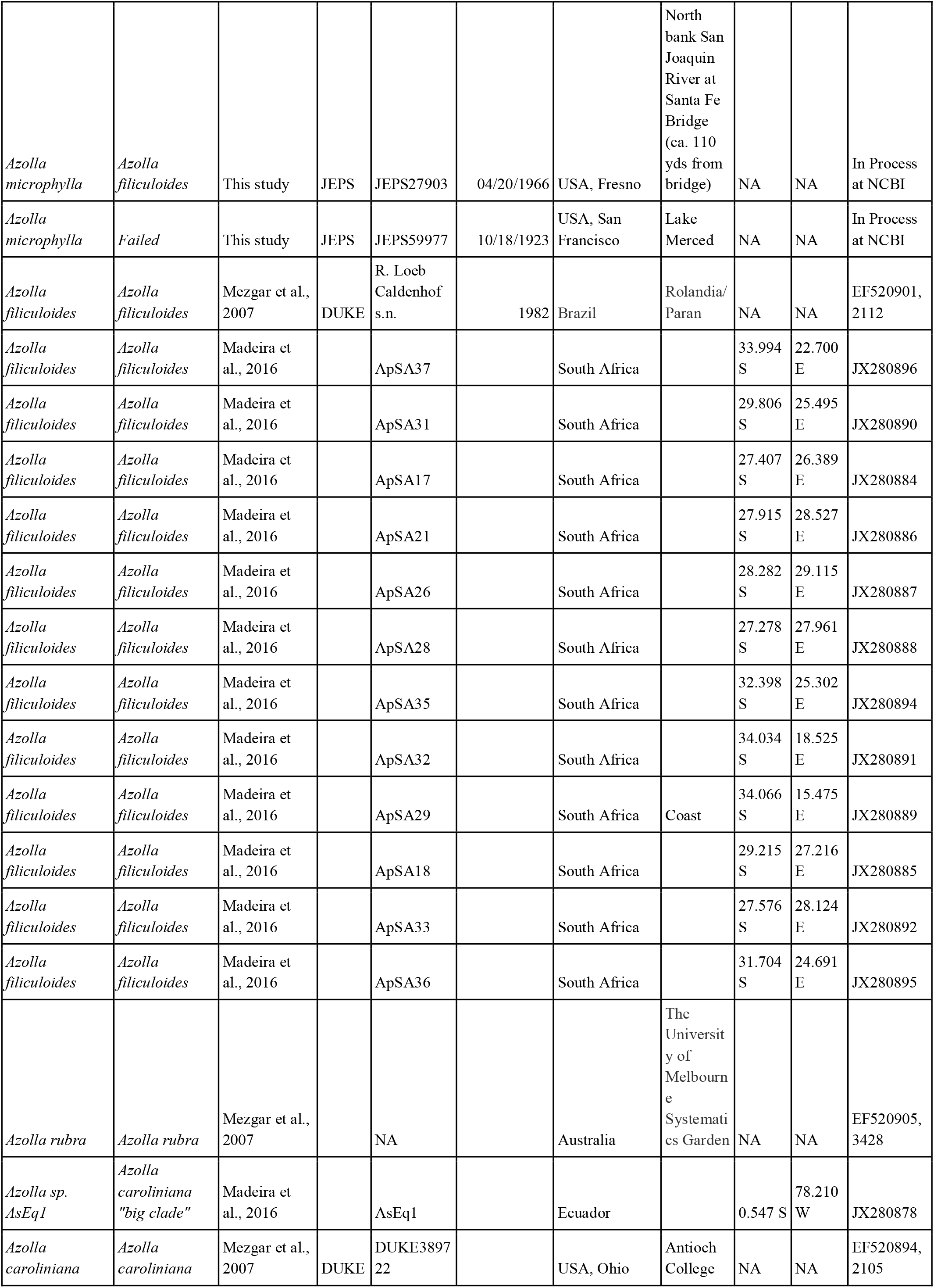
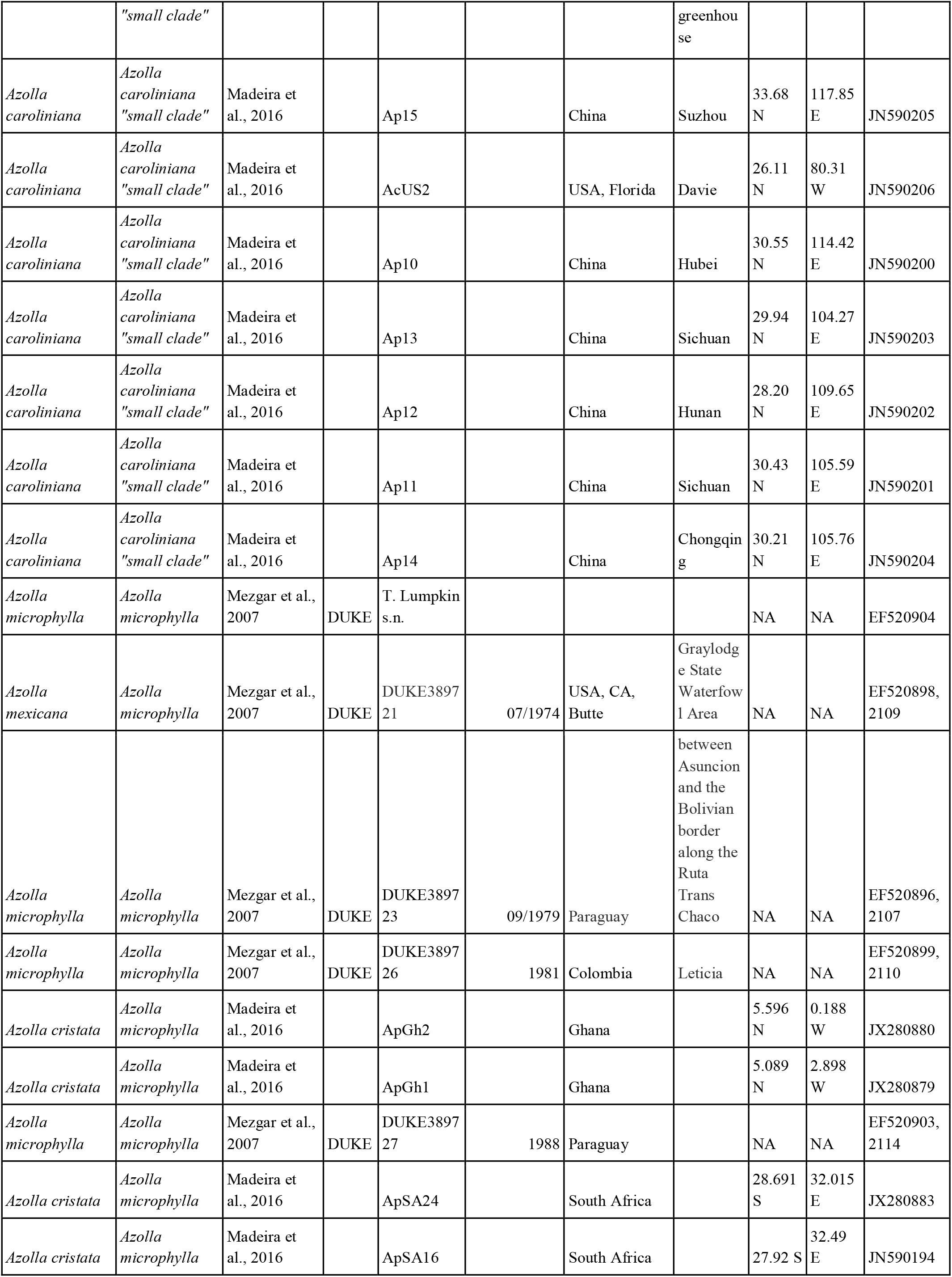
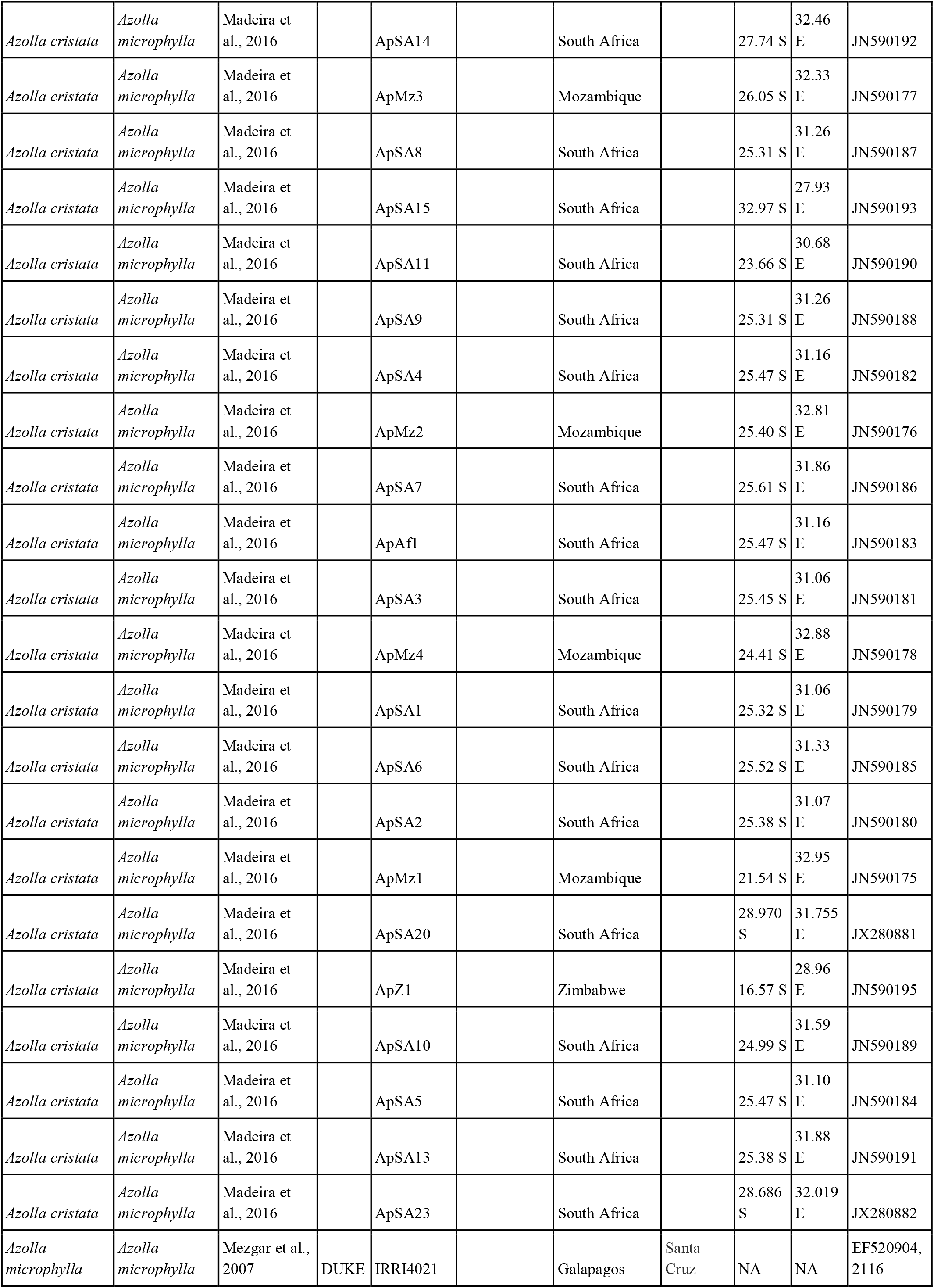
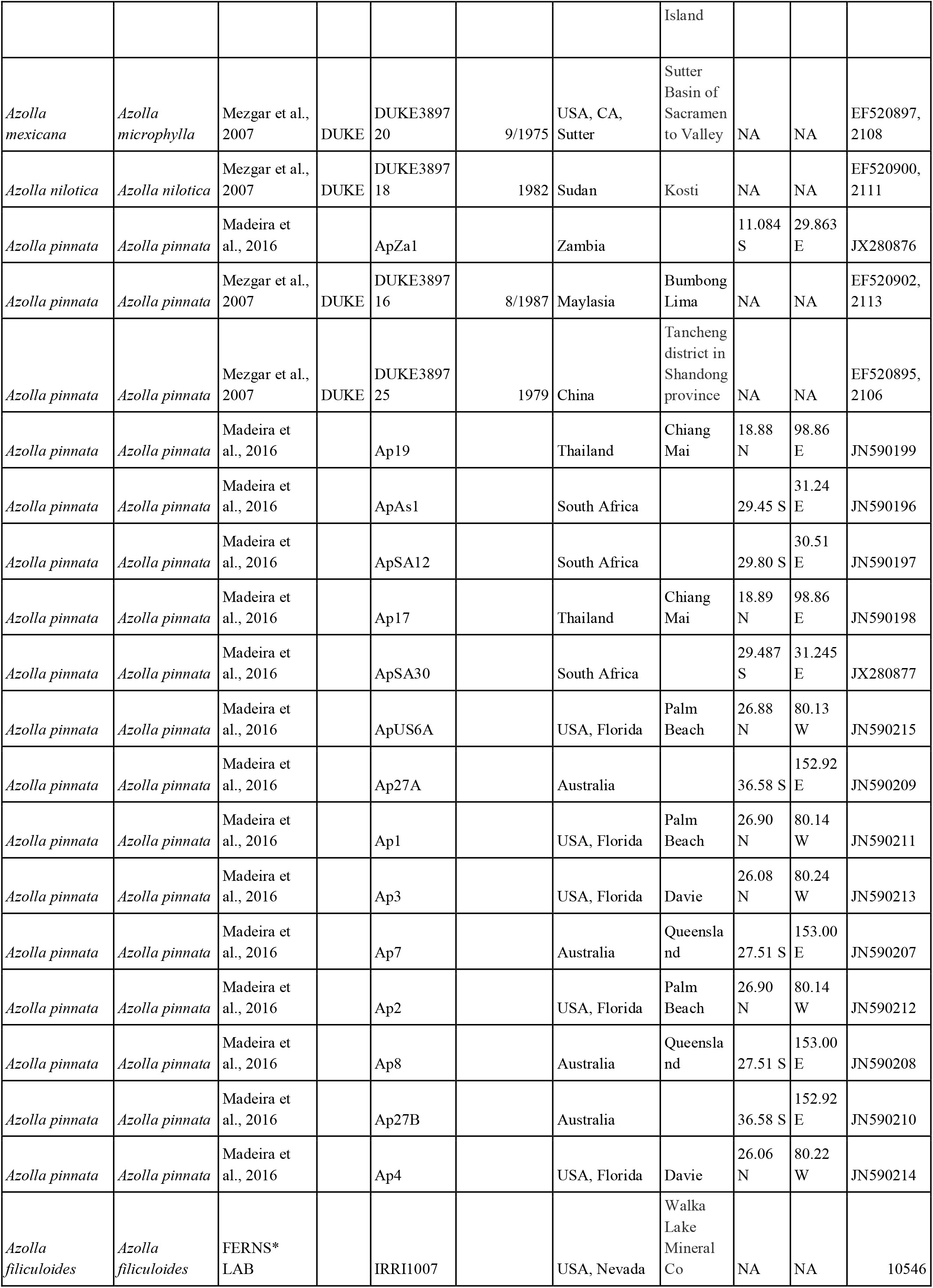
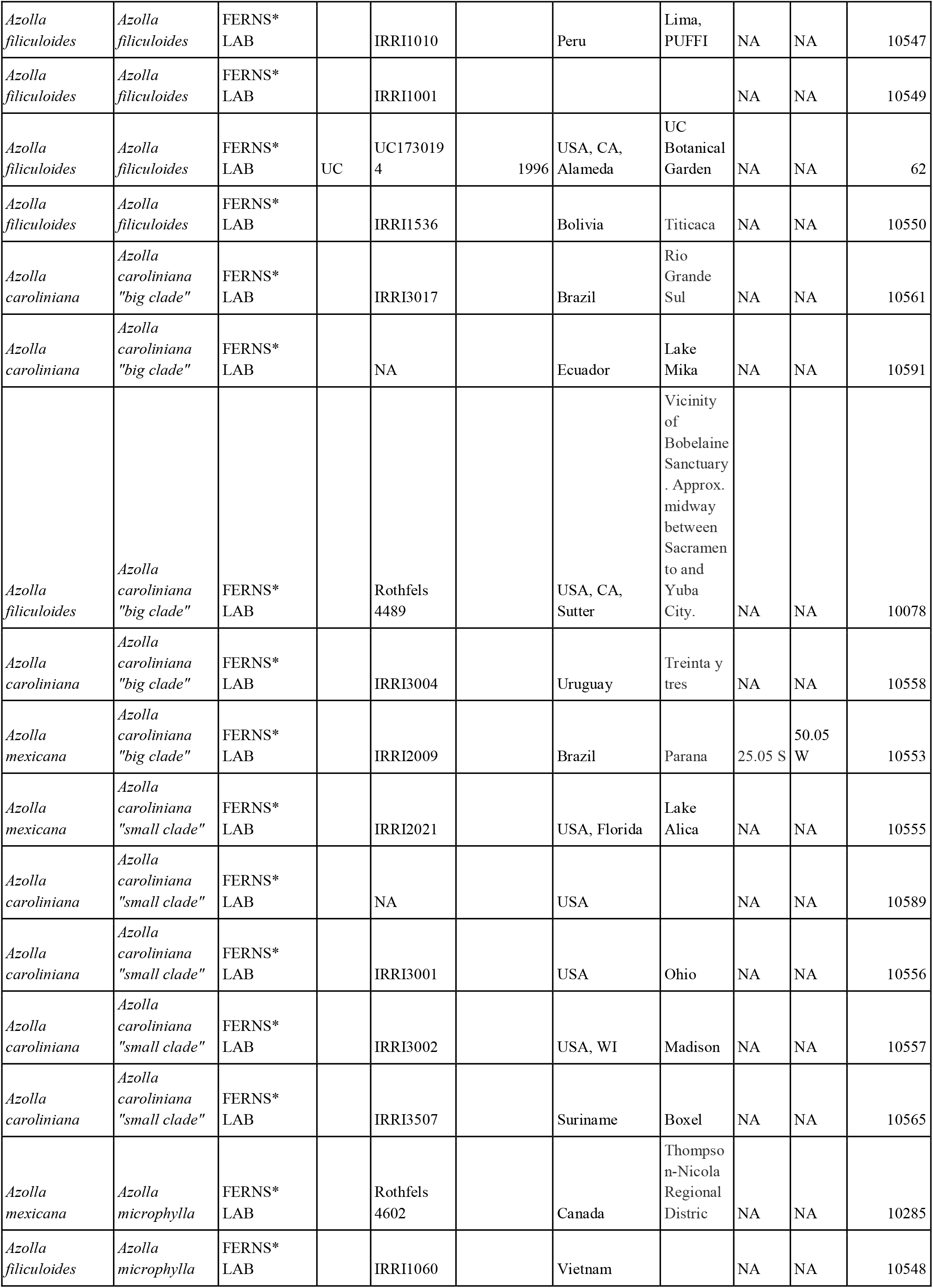
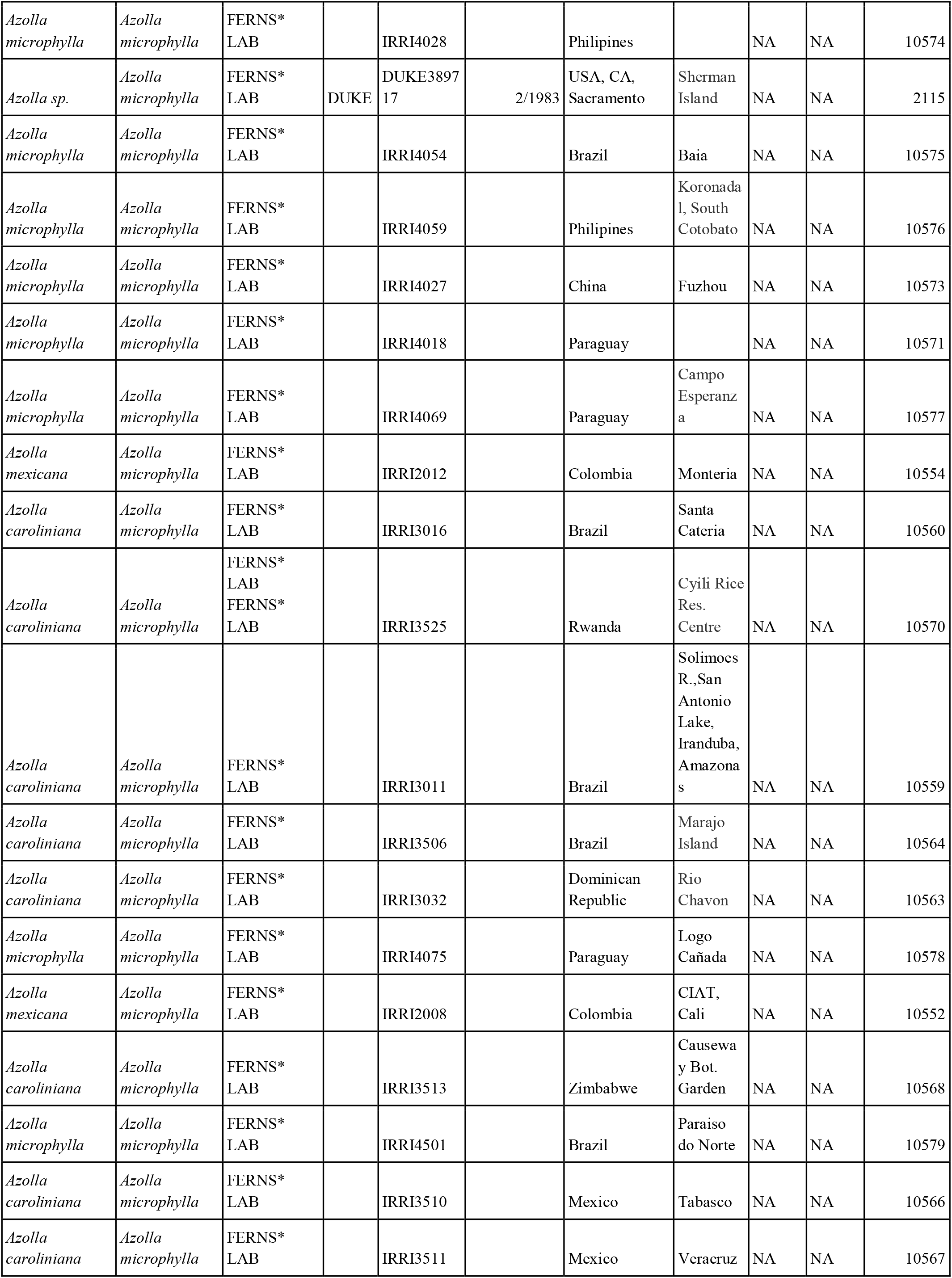
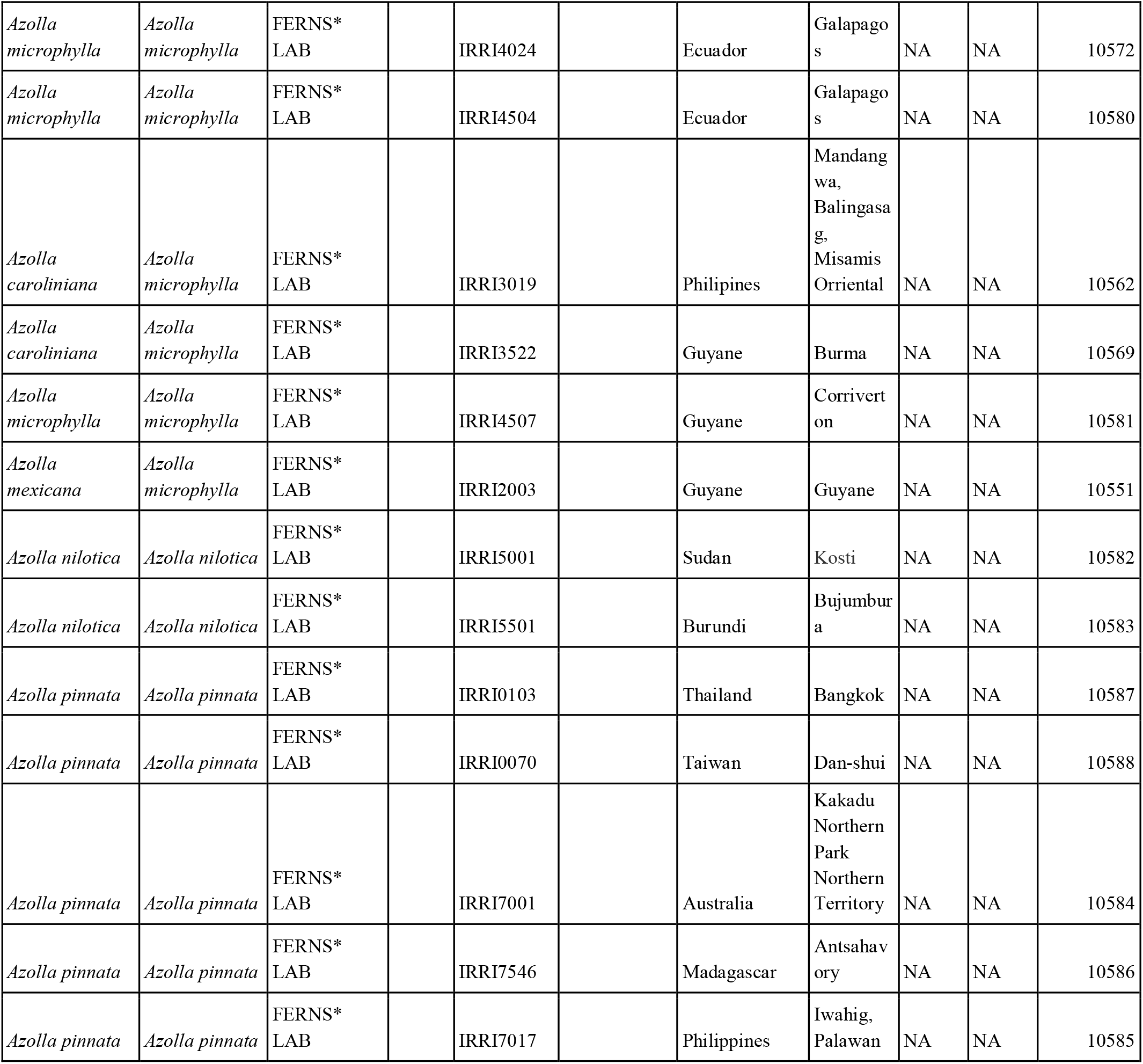
Samples used in this study.

### Morphological examination

We reexamined *Azolla* specimens collected in 2021 to assess morphological differences among the three taxa identified in Song et al. (2025b). We examined the following FERNS* Lab Database vouchers—“*A. caroliniana*” small clade: 15020, 15021, 15022, 15024, 15046, 14970, 14969, 15069, 6859; “*A. caroliniana*” big clade: 14979, 14980, 14981, 14982, 14979, 14980, 14981, 14982, 14983, 14984, 14985, 14987, 14988, 14989, 14990, 14991, 14992, 14993, 14994, 14995, 14996, 14997, 14998, 14999, 15000, 15001, 15002, 15003, 15005, 15006, 15007, 15008, 15009, 15010, 15011, 15012, 15013, 15014, 15015, 15016, 15017, 15018, 15019, 15025, 15026, 15027, 15028, 15029, 15030, 15031, 15032, 15033, 15034, 15035, 15036, 15037, 15038, 15039, 15042, 15043, 15044, 15045, 15047, 15050, 15051, 15052, 15053, 15054, 15056, 15057, 15058, 15059, 15060, 15061, 15071, 15072, 15073, 15074, 10078, 15062; and *A. filiculoides*: 14971, 14972, 6859, 14968, 14976, 15063.

### DNA extraction from herbarium specimens

DNA was extracted in a PCR-product-free laboratory using a DNEasy Plant Mini Kit (Qiagen, Redwood City, CA) following the manufacturers’ standard protocols with two modifications: initial grinding with a micropestle and extension of the final elution step to 10 minutes at 50 °C. Final DNA concentrations were measured using a Qubit fluorometer.

### PCR process

PCR reactions were prepared in a PCR-product-free laboratory using 10 µL water, 7 µL master mix (Invitrogen Platinum), 1 µL of a premixed forward and reverse primer solution, and 2 µL DNA extract template. The following primers were used: TRNG1F GCGGGTATAGTTTAGTGGTAA (Nagalingum et al., 2007), TRNR22R CTATCCATTAGACGATGGACG (Nagalingum et al., 2007), TrnG43F1 GCCGGAATCGAACCCGCATCA (Madeira et al., 2016), TrnG63R TTGCTTMTAYGACTCGGTG (Madeira et al., 2016), rbcLF ATGTCACCACAAACAGAGACTAAAGC (Kress and Ericson, 2007), rbcLE TGATCTCCACCAGACAKACG (Palmieri et al., 2009).

The thermal cycler program was set as follows: 94 °C for 5 min, 40 cycles of 94 °C for 30 s, 45 °C for 30 s, and 72 °C for 60 s, then 72 °C 5 min, and a hold at 4 °C. The annealing temperature was increased to 53 °C for some samples with multiple bands. PCR success was determined using gel electrophoresis. For recently collected *Azolla* samples, we either performed direct PCR or PCR on extracted DNA using the Phire Plant Direct PCR Kit (Thermo Scientific, Waltham, MA) with the same primers and thermal cycler program.

### Sequencing and phylogenetic inference

We sent DNA to Plasmidsaurus Inc. (Watterson Park, Kentucky, USA) for PCR amplicon sequencing with PCR cleanup. For samples where PCR was unsuccessful but DNA extraction was successful, we performed shotgun metagenomic sequencing through Plasmidsaurus (Illumina 2 x 150 bp paired-end reads). Raw reads were processed using Trimmomatic v0.39 (Bolger et al., 2014) with the default parameters for adapter removal and quality trimming. Trimmed reads were assembled using Megahit v1.2.9 (Li et al., 2015) with default parameters on the Galaxy server v26.1.rc1 (The Galaxy Community, 2026).

We performed preliminary BLASTn searches of the sequenced amplicons against the GenBank database to identify *Azolla* species. To recover the *trnG-trnR* and *rbcL* markers from the metagenomic assemblies, we used BLASTn (Altschul et al., 1990) to locate and extract the corresponding sequences. Both amplicons and extracted markers were then analyzed using BLASTn against the NCBI database.

We retrieved publicly available *Azolla trnG-trnR* sequences to create a combined alignment of 260 samples (Table 1). Sequences were aligned using Mafft v7.526 (Katoh & Standley 2013), with default parameters, and the alignment was manually checked in Aliview (Larsson, 2014). A maximum likelihood phylogeny was inferred using IQ-TREE 2 under the GTR model with 1000 bootstrap (UFBoot) replicates (Minh et al., 2020). The alignment, which can also be used by other botanists to identify their *Azolla* specimens, is available at https://michaelsongagradstudent.github.io/azolla/

## Results

PCR amplification was successful for only two of the historical specimens: *trnG-trnR* was amplified from JEPS27903 and JEPS59976, and *rbcL* was successfully amplified from JEPS27903. We recovered partial sequences of both *trnG-trnR* and *rbcL* from JEPS35135 using metagenomic sequencing, but this approach was unsuccessful with JEPS59977. BLASTtn searches identified JEPS27903 and JEPS59976 as *Azolla filiculoides,* rather than *A. microphylla* as was previously determined from reproductive morphology. In contrast, BLASTn identified JEPS35135 as *A. microphylla,* consistent with its existing determination.

The phylogeny inferred from the *trnG-trnR* alignment supported the BLASTn identification of JEPS59976 as *A. filiculoides.* However, JEPS27903 formed a polytomy with *A. microphylla* and the small clade of *A. caroliniana,* potentially due to its partial sequence (Figure 1). Of the 21 samples from our expanded California sampling, 20 were nested within the *A. caroliniana “*big clade” (Song et al., 2025b,c), and one from Mono County was identified as *A. filiculoides* (Figure 1). None was *A. microphylla/mexicana.* Sequenced reads have been made publicly available on NCBI GenBank (Table 1).

**Figure 1:**
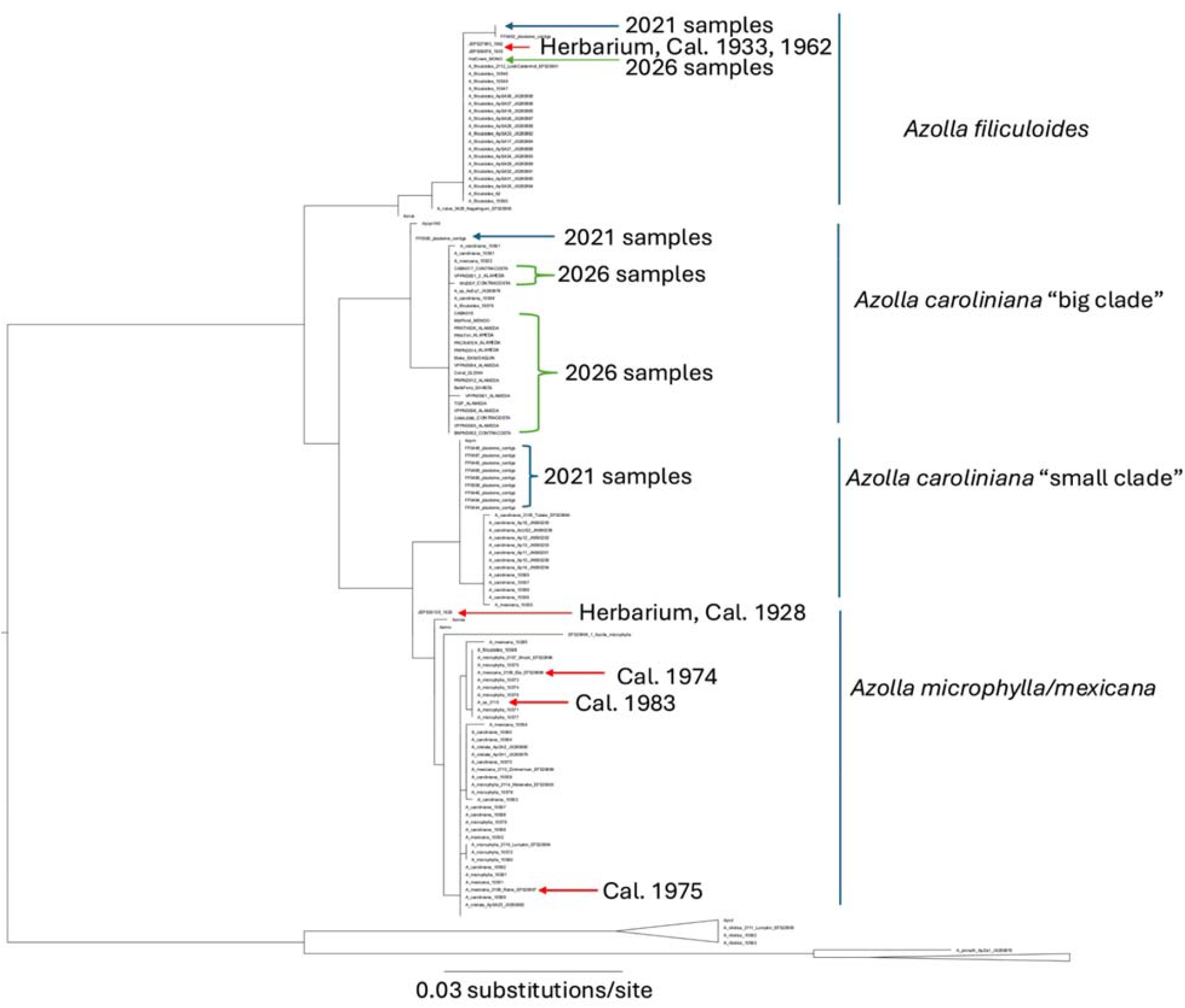
Maximum likelihood phylogeny inferred from *trnG-trnR* sequences. The 2021 samples of *A. filiculoides* and the *A. caroliniana* “big clade” were each collapsed into a single representative tip for readability.

We observed morphological differences between the two “*A. caroliniana”* clades (Table 2). Plants from the “small clade” typically had a leaf fringe 2–3 cells wide near the base and 1–3 cells wide near the apex, as well as conspicuous papillae that appeared to be single-celled. In contrast, plants from the “big clade” typically had a leaf fringe 3–4 cells near the base and 2–3 cells wide near the apex, with variable papillae that were either absent or composed of one or two cells.

**TABLE 2.**
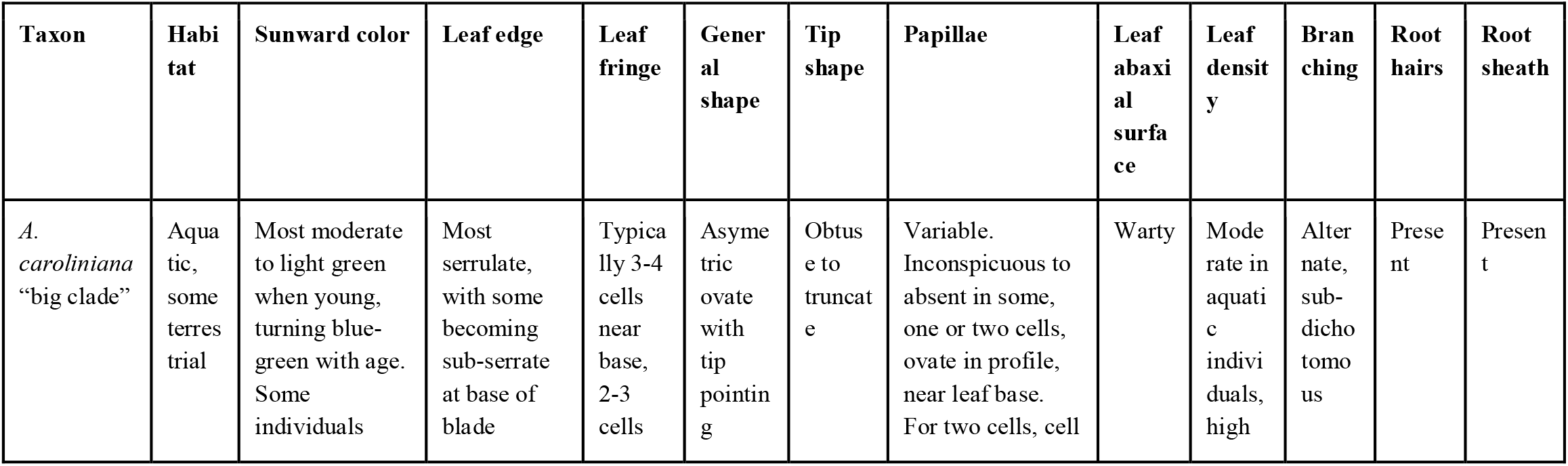
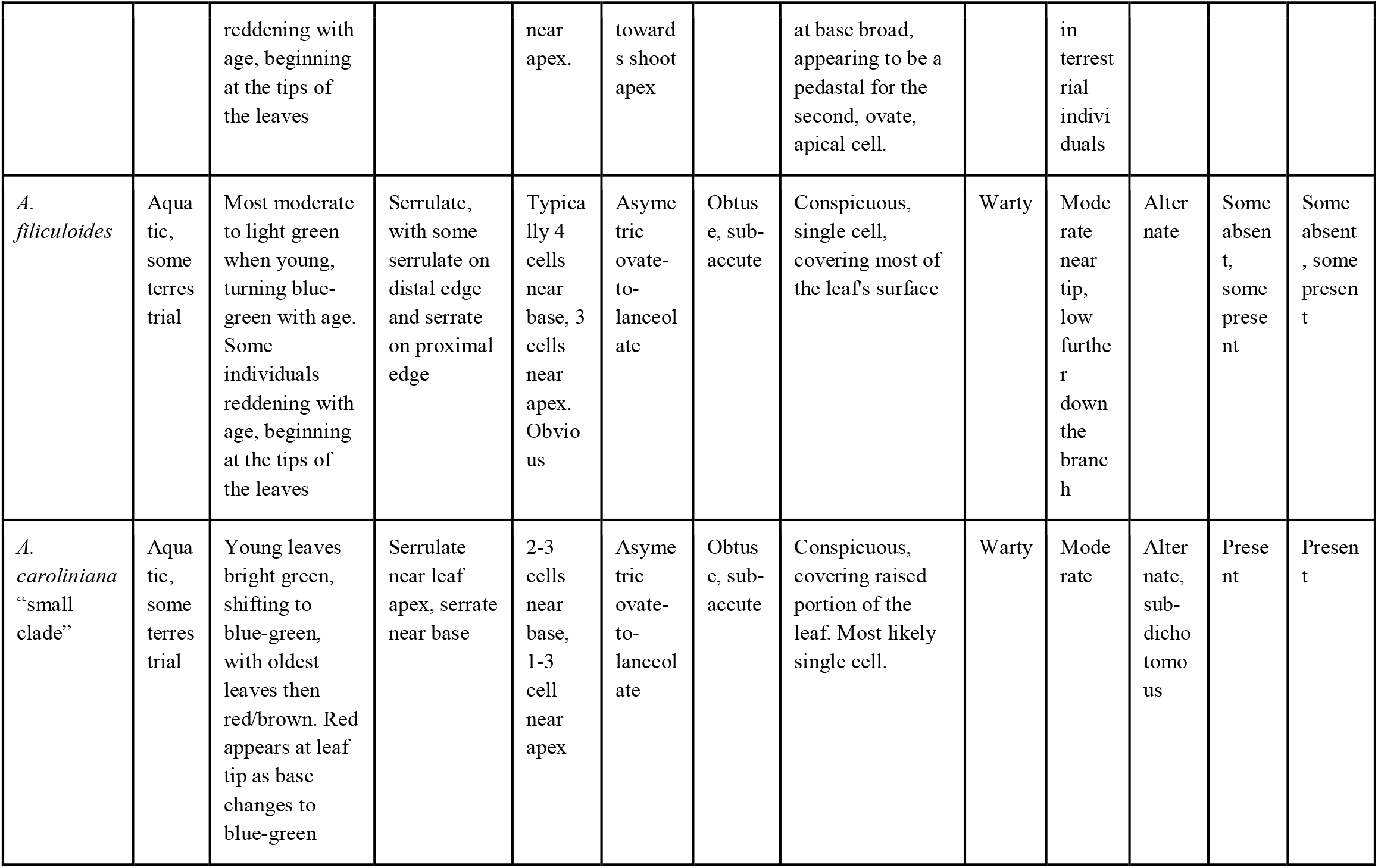
Summary of Azolla morphology examined in this study.

## Discussion

We set out to address three main questions concerning the identification and diversity of California *Azolla*: (1) Can vegetative or reproductive morphological traits reliably identify *Azolla*? (2) Can herbarium specimens yield phylogenetically informative data for *Azolla*? (3) Has there been turnover in California’s *Azolla* diversity over the past several decades? Below, we address these questions and highlight potential directions for future research given our findings.

### Utility of vegetative and reproductive morphological characters for *Azolla* species identification

JEPS59976 and JEPS27903, both previously identified as *Azolla microphylla,* were identified as *A. filiculoides* in our molecular analyses (Figure 1). This is consistent with our findings from Song et al. (2025b,c), in which two samples collected from Kern County in 2023 were also identified as *A. filiculoides*. In contrast, JEPS35135, previously identified as *A. microphylla*, was also placed within the *A. microphylla* clade based on molecular data. Another specimen, DUKE389717, was annotated as originally being identified as *A. filiculoides* and subsequently revised to Azolla sp. following examination of the megasporocarp perine ultrastructure. However, our phylogeny places this specimen within the *A. microphylla* clade. These findings highlight the limitations of morphological characters for identifying *Azolla* and suggest that DNA barcoding can improve the reliability of species determinations. Furthermore, given that local *Azolla* diversity may be underrecognized, identification keys should encompass all species in the genus rather than only those currently recognized in a particular region.

### Value of herbarium specimens and metagenomic sequencing for *Azolla* identification

Our results demonstrate that phylogenetically informative molecular data can be obtained from *Azolla* herbarium specimens that are over 100 years old. We successfully retrieved sequence data from three of the four historical specimens. These results also highlight the utility of shotgun metagenomic sequencing for recovering molecular data from specimens for which traditional PCR is unsuccessful. At the time of this study, PCR amplicon sequencing cost approximately $20 USD per sample, compared with $90 USD per sample for shotgun metagenomic sequencing. Thus, shotgun metagenomic sequencing represents an economically feasible approach for obtaining useful molecular data even from challenging historical specimens.

Relationships among *A. microphylla, A. mexicana,* and *A. carolinana*

Evrard and Van Hove (2004) concluded that morphological characters cannot reliably distinguish the historically circumscribed taxa within the *microphylla-mexicana-caroliniana* clade from morphological traits and propose treating them as one species: *A. cristata*. Reid et al. (2006) and Metzgar et al. (2007) found, using molecular data, that *A. caroliniana* is monophyletic and distinct from *A. microphylla* and *A. mexicana*, but that *A. microphylla* and *A. mexicana* are conspecific. However, genomic studies such as Li et al., (2018) have recognized three species within this group. Song et al. (2025a,b,c) identified two distinct, well-supported clades in California. Each of these clades included specimens identified as *A. caroliniana,* and one of the clades is more closely related to *A. microphylla* and *A. mexicana* than to the other *A. caroliniana* clade. In other words, multiple studies have found support for multiple clades within a broadly defined *A. caroliniana—*and, potentially, for the recognition of three species within the *microphylla-mexicana-caroliniana* complex—but the specific identity of these clades depends on verifying the identity of previously sequenced samples across studies.

The published plastome for *A. microphylla* (MF177089, T. Lumpkin, Ecuador) and the published plastome for a specimen identified as *A. mexicana* (MF177088, D. Rains, USA) are in the same clade as specimens of *A. microphylla* from Ecuador (EF520904, Lumpkin s.n.) and Paraguay (EF520896, Shock s.n. and EF520903, International Rice Research Institute (IRRI): MI4075); *A. mexicana* from Sutter County, CA (EF520897), Butte County, CA (EF520898, IRRI: ME2001), and Colombia (EF520899, IRRI:ME2016); samples of *A. microphylla* and *A. mexicana* from Metzgar et al. (2007); and all of the African *A. cristata* samples from Madeira et al., (2016). This clade includes specimens from South, Central, and North America, as well as Southeast and East Asia, including historical California collections. The genome sequences for *A. mexicana* from Li et al., (2018) are from the same IRRI line (ME2001) as Metzgar et al., (2007) from Gray Lodge, Butte County, California collected by D.W. Rains in 1978. The *A. microphylla* genome from Li et al., (2018) was sequenced from IRRI line MI4021, which is probably the same as the *A. microphylla* from Mezgar et al., (2007). That specimen is also from Ecuador and was also collected by T.A. Lumpkin, although no IRRI number was provided. Evrard and Van Hove (2004) note that the type specimen of *A. microphylla* was collected in California by Chamisso and “disposed of” by Martius and that later descriptions of *A. microphylla* by Mettenius were based on specimens from Cuba. The destruction of the type specimen is notable and further complicates future taxonomic revisions. *Azolla microphylla* was described by Kaulfuss in 1824 and *A. mexicana* was described by Schlechtendal and Chamisso in 1830, so the name *A. microphylla* should take precedence over *A. mexicana*, as has been previously recognized (Evrard and Van Hove, 2004; Smith and Murdock, 2012a,b).

Replacement of historic *A. microphylla* with recent introduction of *A. caroliniana*

Despite thorough and repeated sampling across California, we have yet to verify the continued presence of *A. microphylla* from California. At the same time, we confirm its historic occurrence in California, and recent evidence of widespread populations of *A. caroliniana*. These results provide evidence of a recent introduction of two different *Azolla* taxa into California. (Figure 1).

In the dataset we pulled from the publicly available resources, we found three Californian collections of *Azolla* that come out in our phylogeny as *A. microphylla* —collected in the 1970s and 1980s from Butte County, Sacramento County, and Sutter County. DUKE389721 (A_mexicana_2109_Ela_EF520898 in Figure 1) was collected by Steve Ela in July 1974 at the Graylodge State Waterfowl Area, Butte County, CA. DUKE389717 (A_sp_2115 in Figure 1) was collected in February 1983 by Steve Talley at Sherman Island/Expt. 1052 in agricultural Sacramento County, CA. DUKE389720 (A_mexicana_2108_Rains_EF520897 in Figure 1) was collected in September 1975 in the Sutter Basin of Sacramento Valley, Sutter County, CA in a saline intruded area at an elevation of 10 meters. While we sampled Sacramento and Sutter Counties extensively in 2021 and did not find *A. microphylla*, Butte County should be prioritized for future sampling to confirm the absence or presence of *A. microphylla* in California.

Two potential species of *A. caroliniana*

Samples identified as *A. caroliniana* come out in two distinct clades. While the two clades inferred from plastid markers would render it paraphyletic as currently circumscribed, our previous study (Song et al., 2025a) that used nuclear markers found that these two clades are sister to each other, yet still distinct. What, then, is *A. caroliniana*? Let us first turn our attention to the smaller clade of *A. caroliniana* (Figure 1). This group of mostly California samples with one North Carolina sample (FF0490) is nested within a clade of *A. caroliniana* that includes samples such as the Ohio specimen (EF520894; IRRI3001; FernLabDatabase:10556) from Metzgar et al. (2007) and Reid et al. (2006). Reid et al. (2006) also included a sample from Wisconsin (IRRI CA3002) and the Philippines (IRRI PI0001). This Wisconsin specimen is also found within this first *A. caroliniana* clade that also includes collections from Florida and China. This clade is probably what has historically been considered *A. caroliniana*, i.e. the plant native to the Eastern United States. It is also of note that Pereira et al. (2011) found two groups of *A. caroliniana* within their dendrogram of vegetative character similarities that correspond to the two clades of *A. caroliniana* that we found: one group corresponds to the big clade (IRRI 3017, Brazil) and one to the small clade (IRRI 3001, 3002, USA). However, this second group of Pereira et al. (2011) also includes samples that we identify as *A. microphylla* (IRRI 3513, 3525, Africa) and not all of the samples were shared across studies. Reid et al. (2006) suggest that *A. caroliniana* needs a lectotype or neotype, as the original type is probably conspecific with *A. filiculoides*.

Reid et al., (2006) recovered *A. caroliniana* as monophyletic using plastid markers, potentially because their sampling did not include South American specimens. Additionally, none of the published whole genomes and whole plastomes of *A. caroliniana* come from the east coast of North America. Li et al. (2018) published two *A. caroliniana* genomes. The first genome comes from a sample of IRRI 3017 from Rio Grande do Sul in Brazil originally collected by I. Watanabe in 1987. This may correspond to the “Acar140” plastome on Genbank (MF177092) which does not have an IRRI number but was also collected by Watanabe from Brazil. The second genome was sequenced from a sample of IRRI 3004 collected from Treinta y tres in Uruguay by D.W. Rains in 1982. This likewise may correspond to the “Acar145” plastome on NCBI (MF177093), which has the same location and collector, but no IRRI number. These plastome sequences are nearly identical to the big clade of *A. caroliniana* from Song et al. (2025b,c) and are closely related to other collections from Uruguay, Brazil, Ecuador, and interestingly to a collection from Sutter County in California by Rothfels in 2014. This clade of *A. caroliniana* is sister to a clade of the other clade of *A. caroliniana* plus *A. mexicana-microphylla*. These relationships suggest that three species may be represented in this group, although further taxonomic work is needed to establish appropriate nomenclature of this South American lineage now widespread in California.

Another complication that this phylogeny reveals is that the ranges of the three species overlap more substantially than previously appreciated. Notably, all three lineages have been at some point moved all around the world well beyond their hypothesized native ranges. It remains to be determined how these three essentially identical species with overlapping niches maintain species boundaries. Nonetheless, there are a few morphological characters that may help distinguish the two putatively different taxa currently treated as *A. caroliniana*. More work will need to be done to officially assign proper binomial names to these clades given the extreme nomenclatural problems of the group, if they are to be recognized at the level of species (Evrard and Van Hove, 2004). Evrard and Van Hove (2004) propose recognizing only two species of *Azolla* in North America, *A. filiculoides* and *A. cristata* (syn. *A. mexicana*, *A. caroliniana*, *A. microphylla*). *Azolla filiculoides* has unicellular trichomes on the upper leaf lobe whereas the trichomes of *A. cristata* are multicellular. However, one of the *A. caroliniana* clades examined in our study frequently had unicellular trichomes, despite its phylogenetic placement outside *A. filiculoides*. Therefore, we are in agreement with Bates (1980) and Tan et al., (1986) that the leaf trichome may not be a particularly useful character. More work is needed to determine whether combinations of morphological characters can reliably distinguish *Azolla* species.

## Acknowledgments

We would like to thank the Jepson and University of California Herbraria (JEPS, UC) for providing samples, especially Sonia Nosratinia and Ana Penny. For assistance in collecting or providing access to field samples of *Azolla*, we thank Adina Merenlender and Evelyn George (University of California Hopland Research and Extension Center), Tammy Lim and Edward Culver (East Bay Regional Parks District), Jessica Purificato (East Bay Municipal Utility District), Jack Respet, Nicholas Masi, Paige McCreadie, Gracelyn Ham, and Jordayn Eichhorn (University of Colorado, Boulder). For providing research facilities and logistical support, we would like to thank Zac Harlow and Zac Tuthill (UC Berkeley Blue Oaks Ranch Reserve). Field collections were funded, in part, by the Department of Defense Strategic Environmental Research and Development Program (SERDP Project RC24-4111: 101897) and the CU Boulder EBIO Graduate Student Research Award. This work was supported in part by the National Science Foundation Research and Mentoring for Postbaccalaureates in Biological Sciences (RaMP) program (Award No. 2216550) and the College of Science and Engineering at San Francisco State University (SFSU). We thank the SFSU Genomics and Transcriptomics/Analysis Core for providing facilities and support for molecular analyses.

## Notes

### Competing Interest Statement

The authors have declared no competing interest.

